# Taxon-specific differences in C and N cycling and metabolic activity of intertidal organisms: Part B – Long-term processes

**DOI:** 10.64898/2026.09.07.749936

**Authors:** Tanja Stratmann, Dick van Oevelen, Marcel T. J. van der Meer

**Author notes:** **Correspondence:** Tanja Stratmann.

## Abstract

Tidal flats cover less than 1% of the surface of our planet, but they provide important ecosystem functions and services, like carbon sequestration and nutrient (re-)mineralization, primary, secondary, and food production, and habitats for a variety of species. Despite their importance, they are threatened by environmental stressors and anthropogenic impacts. Hence, a better understanding of the interaction between macrozoobenthos, their metabolic activity, and their contribution to carbon and nitrogen cycling is important. Therefore, we conducted *in-situ* pulse-chase tracer experiments at a Pacific oyster (*Magallana gigas*) reef in the Eastern Scheldt (Southwest Dutch Delta, Netherlands) in summer and autumn 2020. During these experiments, we provided the submerged benthic community with ^13^C and ^15^N-enriched bacterioplankton and incubated it in seawater enriched in 1% deuterium oxide (^2^H_2_O) for one tidal cycle (∼9 h), before the benthic community was kept at *in situ* conditions for another 14 d. Our aim was (1) to assess seasonal variations in ^13^C and ^15^N processing by macrozoobenthos, (2) to evaluate potential life-stage dependent differences in ^2^H uptake – a proxy for metabolic activity, and (3) to identify the macrozoobenthos taxon which had the highest metabolic turnover rate (*MTR*). In summer, most bacterioplankton-derived ^13^C- and ^15^N was taken up by sponges (*Halichondria panicea*), limpets (*Crepidula fornicata*), and clams (*Ruditapes philippinarum*), whereas in autumn, *H. panicea, C. fornicata* and the sponge *Hymeniacidon perlevis* dominated ^13^C and ^15^N incorporation. Statistically significant differences in ^2^H uptake between life stages (juvenile/ small vs. adult) was detected in Malacostraca (i.e., *Rhithropanopeus harrisii*), but not in Polychaeta or Bivalvia. Therefore, this difference between taxonomic classes might be linked to mold-intermolt periods. Seasonal differences in *MTR* of *H. panicea* and *M. gigas* (i.e., the species with the highest *MTR*s in summer and autumn) might be linked to seasonal environmental variability, such as temperature and food availability.

## Introduction

Tidal flats, i.e., coastal areas that are submerged during high tide and exposed to air during low tide, like mud, sand, and rock flats, cover at least 0.025% of the surface of our planet (Murray et al., 2019; Chapman et al., 2025). Most tidal flats are located in Asia, North America, and South America (Murray et al., 2019). Europe (excluding Russia) has about 10,252 km^2^ of tidal flats (Hill et al., 2021) that sporadically host non-native Pacific oyster *Magallana gigas* reefs. For instance, in the Dutch Wadden Sea and in the Eastern Scheldt (Oosterschelde, Dutch Delta, South-western North Sea) 0.16% (de Jonge and Boddeke, 1993; Fey et al., 2010) to 6.72% (Smaal et al., 2009) of the tidal flats are covered with *M. gigas* reefs. In the eastern part of the Eastern Scheldt, these oyster beds even spread over 9.12% of the intertidal area (Smaal et al., 2009).

Intertidal mudflats and oyster reefs provide various ecosystem functions, such as habitat (De Santiago et al., 2019; Chan et al., 2022), carbon (C) and nutrient cycling (Dame et al., 1989; Middelburg et al., 1995; Kellogg et al., 2013; Westbrook et al., 2019; Rios-Yunes et al., 2023a). Other ecosystem functions are primary and secondary production (Sprung, 1994; Brotas and Catarino, 1995), biodiversity (Guy et al., 2018; Bijleveld et al., 2025; Martin et al., 2025), sediment resuspension and trapping (Carey, 1983; Reidenbach et al., 2013), or wave attenuation (Morris et al., 2021; Salatin et al., 2022). Additionally, these mudflats and oyster reefs provide ecosystem services, like ‘regulating services’ (Grabowski and Peterson, 2007; Passarelli et al., 2018; Chen and Lee, 2022; Giglio et al., 2024), ‘provisioning services’ and ‘cultural services’ (Grabowski and Peterson, 2007; Passarelli et al., 2018; Thomas et al., 2022; Giglio et al., 2024).

Many of these ecosystem functions and services are linked to the presence and activity of macrozoobenthos (Herman et al., 1999; Thrush et al., 2006). However, this macrozoobenthos is particularly vulnerable to environmental and anthropogenic stressors, such as atmospheric and marine heatwaves (Raymond et al., 2024; Hamer et al., 2026), storms (Martinez et al., 2022; Liu et al., 2023), habitat loss due to rapid expansion of the human population (Klein et al., 2026) and shoreline retreat (Zhang et al., 2019), pollution with chemicals (Mazik and Elliott, 2000), heavy metals (Muhaya et al., 1997; Li et al., 2024), oil (Nwipie et al., 2019), and microplastics (Vermeiren et al., 2023; Kneel et al., 2024). Therefore, it is important to understand how macrozoobenthos interacts, how it (re-) cycles C and nitrogen (N), and how metabolically active it is. Here, metabolic activity is the “chemical changes […] in living animals […] as a result of [their] metabolism” (Park, 2012) and it is measured as the uptake of deuterium (^2^H) (Stratmann et al., 2025a, 2026).

Short-term (12 h) *ex-situ* incubation experiments with *M. gigas*, its epibiont, the sponge *Halichondria* (*Halichondria*) *panicea*, and further associated (smaller) fauna from the Eastern Scheldt showed that the crabs *Carcinus maenas* and *Eriocheir sinensis, H. panicea*, and the snail *Littorina littorea* had the highest ^2^H uptake and consequently were the most metabolically active species (Stratmann et al., 2026). Most of these very metabolically active species had incorporated bacterioplankton-derived ^13^C- and ^15^N via filter and suspension feeding (i.e., primary consumer/ direct uptake; *H. panicea*) or via predation upon sponges (i.e., secondary consumer/ indirect uptake; *C. maenas, E. sinensis*), indicating a direct link between feeding and metabolic activity (Stratmann et al., 2026). Besides assessing the latter two activities, it should also be possible to estimate the metabolic turnover rate (*MTR*, % d^-1^) of individual species. Here, *MTR* refers to the organism’s renewal rate of parts of its tissue, organs, and/ or body fluids measured as the change in ^2^H incorporation over time.

The *MTR* can be calculated using data from pulse-chase tracer experiments in which the taxon of interest is incubated in the presence of ^2^H at T_0_ (i.e., the ‘pulse’), and the change in the concentration of incorporated ^2^H is measured over time at T_1_, T_2_, T_3_, etc. (i.e., the ‘chase’). During the ‘pulse’ part in which the taxon is incubated in ^2^H_2_O, the taxon incorporates ^2^H from ambient seawater. After the source of ^2^H_2_O has been removed, the taxon continues to incorporate mainly natural abundance ^1^H from the ambient water, thus diluting the ^2^H enrichment of its tissue. This loss in ^2^H enrichment measured during the ‘chase’ part is directly related to cell division, the build-up of new tissue, and potentially to somatic growth – though the latter still needs to be tested.

In this study, we conducted *in-situ* pulse-chase tracer experiments at intertidal mudflats with oyster reefs with ^13^C- and ^15^N-enriched bacterioplankton as food source (i.e., substrate-dependent stable isotope tracer) and ^2^H-enriched seawater as substrate-independent stable isotope tracer or “substrate-agnostic isotope tracer” (Callaghan et al., 2023). With this experimental design, we aimed (a) to determine the long-term (14 days) processing of bacterioplankton by an intertidal community in summer and autumn, (b) to assess potential differences in metabolic activity depending on the life stage of macrozoobenthos, and (c) to identify the macrozoobenthos species with the highest *MTR*.

## Materials and methods

### Study site

As a semi-enclosed tidal bay in the Dutch Delta, the Eastern Scheldt is 350 km^2^ large and has a tidal range of 2.9 to 3.5 m (Jiang et al., 2019a). It has a tidal exchange of ∼20,000 m^3^ s^-1^ (Jiang et al., 2019b) and its seawater is turned over every 90 to 120 days (Jiang et al., 2019b). About 62% of the total area of the Eastern Scheldt is tidal shoals, the intertidal zone (i.e., the zone between mean low water and mean high water) comprises 36% of the area of the bay, and 2% of the area is salt marshes (Louters et al., 1998). In the intertidal zone, 0 to 51% of the sediment is silt (<63 µm) (van der Wal et al., 2010; Daggers et al., 2018; De Borger et al., 2020), whereas in the subtidal, 24 to 63% of the sediment is silt (De Borger et al., 2020). The median grain size is 106 ± 38.6 µm in the intertidal and 126 ± 72.0 µm in the subtidal (Rios-Yunes et al., 2023b).

### Experimental design

*In-situ* experiments (*n* = 6) and the corresponding controls (*n* = 6) were performed in summer (July) and autumn (September – October) 2020 (Supplementary Table 1, Supplementary Figure 1) at an intertidal *M. gigas* reef in the eastern part of the Eastern Scheldt (51.489°N, 4.058°E; Fig. 1). For the controls, specimens with natural abundance stable isotopic composition were collected.

**Figure 1.**
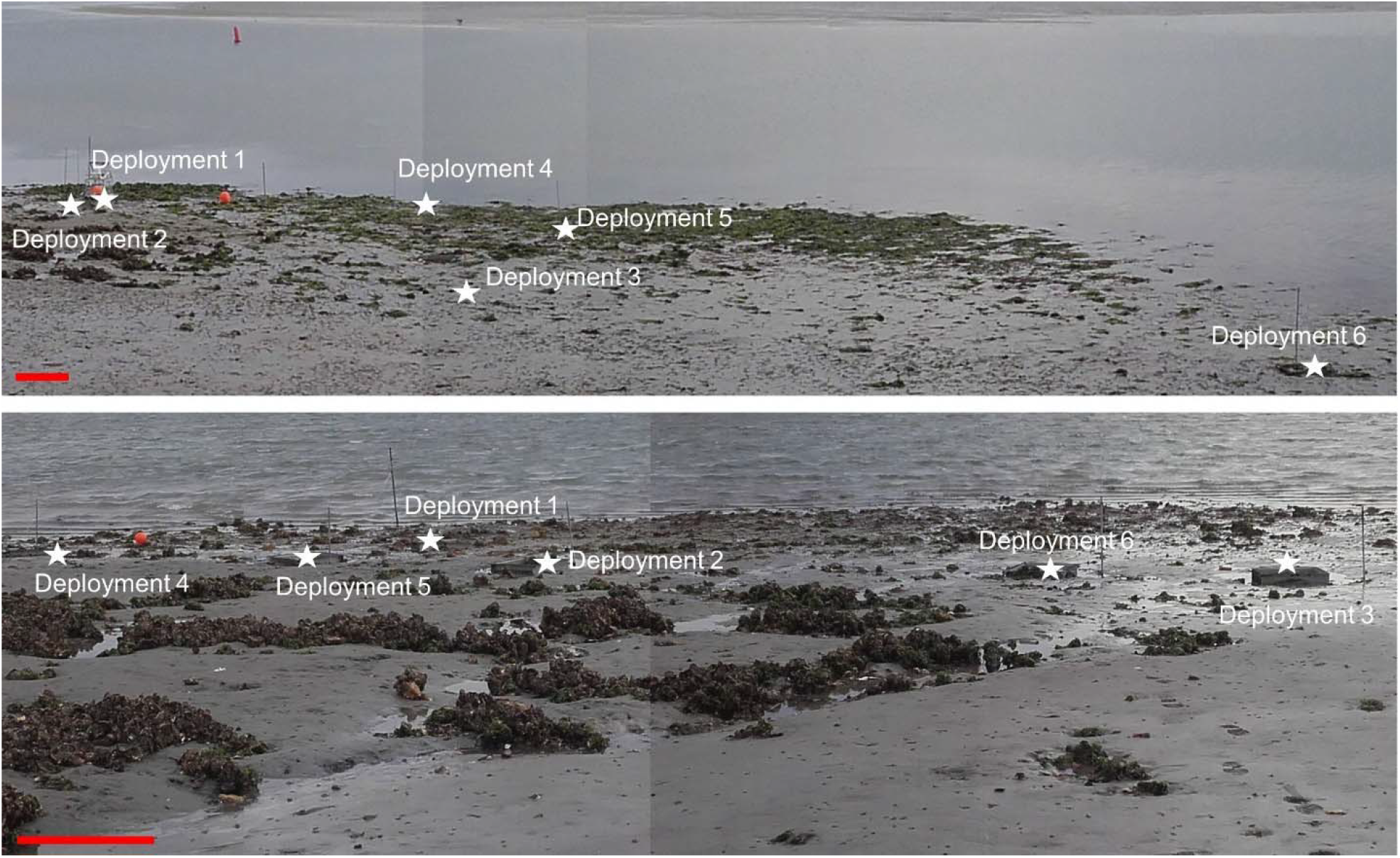
Photomosaic of the experimental sites in the Eastern Scheldt (Dutch Delta, southwestern North Sea) during low tide in (upper panel) summer and (lower panel) autumn 2020. White stars indicate the location of each experimental deployment. The red scale bar represents 1 m.

### Main treatment

During low tide, six 50 × 50 cm stainless steel frames were buried until only 3 – 5 cm of the frames protruded from the sediment (Supplementary Figure 2). These steel frames served as foundation on which the incubation chambers CUBEs (chamber size: 50 × 50 × 50 cm (Stratmann et al., 2018, 2023); Supplementary Figure 2) were placed for the so-called ‘incubation period’ (1 tidal cycle, full submergence: ∼9 h). After the incubation, the metal foundation was kept in place for the so-called ‘turnover period’ (14 d).

Immediately before the start of the ‘incubation period’, the CUBEs were placed on the frames and fixed before the high tide, while the openings (door, port in the ceiling) were kept open so that the seawater could fill the chambers during rising tide. When the water level almost reached the plexiglass ceiling, the doors were closed and fixed. Deuterated water (^2^H_2_O; 1.25 l, 99.9% ^2^H_2_O, Eurisotop, France; equivalent to 1% (v/v) ^2^H-enrichment) and (mean ± std) ∼34.2 ± 13.2 ml ^13^C and ^15^N-enriched concentrated substrate bacteria (Stratmann et al., 2026) (summer 2020: no data; autumn 2020: 29.0 ± 1.59 at% ^13^C/ 47.3 ±1.68 at% ^15^N; 105 ± 55.0 mg ^13^C, 31.1 ± 12.7 mg ^15^N; Supplementary Table 2) were poured through the port into the chamber. Directly afterwards, duplicate water samples for the quantification of inorganic nutrients (ammonia, nitrate, nitrite, phosphate, dissolved silicate) were taken through the sampling port at T_0_ and a water sample was taken from the seawater outside the chamber (T_-1_). The water was filtered through 0.45 µm pore size filters into 6 ml vials and stored frozen (-20ºC; nitrite, ammonia, nitrate, phosphate) or cold (4ºC; dissolved silicate) until analysis. Water samples for the quantification of dissolved inorganic C (DIC) and ^13^C-DIC were taken in 10 ml headspace vials, preserved by adding 10 μl saturated HgCl_2_, and stored at 4°C.

During the ‘incubation period’, water inside the chambers was kept well mixed by a rotating plate inside the chamber (see (Stratmann et al., 2018)). Additional water samples were taken by the CUBES automatically with 30 ml syringes in 1.5 h time intervals (T_1_: 1.5 h after tracer addition, T_2_: 3 h after tracer addition, T_3_: after 4.5 h, T_4_: after 6 h, T_5_: after 7.5 h, T_6_: after 9 h). When the CUBEs re-emerged from the high tide during the next low tide, they were taken off the frames, transported back to the lab (approximately 5 min walking distance), and the water in the syringes was prepared for analysis of inorganic nutrients, (^13^C-) DIC as described above.

After 14 d at the end of the ‘turnover period’, sediment inside the metal frames was sampled as follows: Three plexiglas cores for benthic bacteria, one plexiglas core for total organic carbon (TOC), and one plexiglas core for phytopigments (inner Ø of all cores: 3.5 cm), were pressed into the sediment (Supplementary Figure 3). Additionally, one plexiglass core (inner Ø: 3.5 cm) per season was pressed into the sediment to take samples for grain sizes analyses. After all *H. panicea* and *M. gigas* specimens at the sediment surface were collected by hand, the sediment-filled cores were retrieved, and the remaining sediment was excavated to about 10 cm of depth. This sediment was sieved (500 µm pore size) in the field using seawater from a tidal channel of the Eastern Scheldt. All macrozoobenthos infauna (>500 µm) that remained on the sieves (Supplementary Figure 3) were collected and transported back to the lab.

In the lab, the sediment cores for benthic bacteria, TOC, and phytopigments analyses were sliced in four intervals (0.0 – 0.5 cm, 0.5 – 1 cm, 1 – 3 cm, 3 – 8 cm) and immediately frozen (benthic bacteria, TOC samples: -20°C, phytopigment samples: -80°C). *H. panicea* specimens were washed in pre-filtered seawater (2 μm pore size) and scrapped off any hard substrate (e.g., clinker, shells, parts of roof tiles). Subsequently, all associated fauna growing attached to *H. panicea* as epifauna or living inside *H. panicea* tissue were handpicked and frozen (-20°C). Wet mass (WM) of *H. panicea* was determined and each specimen was frozen individually (-20°C). After washing with pre-filtered seawater, *M. gigas* specimens were opened with oyster knives, the flesh was removed, WM was measured, and it was frozen (-20°C). Any epifauna growing on *M. gigas*’ shells was handpicked and frozen (-20°C), before the shells were discarded. The macrozoobenthos infauna was screened under a stereomicroscope to separate debris, gravel, or empty shells, from living macrozoobenthos which was fixed in borax-buffered 4% formaldehyde.

### Control

In each season, a 50 × 50 cm plastic frame (Supplementary Table 3, Supplementary Figures 2 and 4) was placed six times at the control site near the *in-situ* experimental site during low tide. Each time, one core for TOC, one core for phytopigments, and three cores for benthic bacteria were placed inside the frame as described in “Main treatment”. One additional core per season was taken for grain size analyses. After all cores were retrieved and all *H. panicea* and *M. gigas* specimens inside the frame were collected by hand, the remaining sediment was excavated to 10 cm depth, and sieved at site over a 500 µm pore size sieve.

In the lab, sediment cores, macrozoobenthos samples, *H. panicea* and *M. gigas* specimens were processed and preserved as described in “Main treatment”.

### Sample processing

#### Macrozoobenthos samples

Macrozoobenthos samples were counted and identified to the finest taxonomic level possible (i.e., mostly species/ genus level for adult specimens, higher taxonomic level for juveniles) under a stereomicroscope using (Hayward and Ryland, 1995; Bos et al., 2016) as references. Subsequently, every macrozoobenthos specimen was freeze dried, weighed with shell and without shell (i.e., DM), and ground to powder with pestle and mortar. Organic C (OC) content/ δ^13^C, total nitrogen (TN) content/ δ^15^N, and H content/ δ^2^H of every specimen were measured with an elemental analyzer coupled to an isotope ratio mass spectrometer (EA-IRMS) as described in (Stratmann et al., 2025a).

Stable isotope values of C, N, and H were reported in δ notation relative to Vienna Pee Dee Belemnite (VPDB) for δ^13^C, relative to air for δ^15^N (Fry, 2006), and relative to Standard Mean Ocean Water normalized to Standard Light Antarctic Precipitation (VSMOW-SLAP) for δ^2^H (Coplen, 1995).

#### Sediment samples

Sediment samples were freeze dried, ground with mortar and pestle to powder, and OC content/ δ^13^C and TN content/ δ^15^N were measured with an EA-IRMS as described in (Stratmann et al., 2023, 2025b) (instrument setting is presented in Additional Information A1 of (Stratmann et al., 2025a)).

#### Fatty acid extraction and analysis

Phospholipid-derived fatty acids (PLFAs) were extracted from ∼3 g freeze-dried, fine-ground sediment following the modified Bligh and Dyer extraction method (Bligh and Dyer, 1959; Boschker, 2008) as described in detail in (Stratmann et al., 2023).

After derivatizing the PLFAs to fatty acid methyl esters (FAMEs) using mild alkaline methanolysis, C concentrations of individual FAMEs were measured with a gas chromatograph with a flame ionization detector (GC-FID) using a polar BPX70 column as described in (Stratmann et al., 2025a) (instrument setting is presented in Additional Information A2 of (Stratmann et al., 2025a)). Individual FAME peaks were identified in the chromatograms based on their equivalent chain length (ECL) in relation to ECLs of the internal standards C12:0 FAME and C19:0 FAME, and the omnipresent peak of C16:0 FAME.

The δ^13^C_VPDB_ and δ^2^H_VSMOW-SLAP_ values (‰) of individual FAMEs were measured on a GC-IRMS using a BPX70 column as described in (Stratmann et al., 2025a) (instrument setting is presented in Supplementary Table 4).

#### Inorganic nutrient and dissolved inorganic carbon analyses

Inorganic nutrient (ammonia, nitrite, nitrate, dissolved silicate, phosphate) and DIC concentrations (µmol l^-1^) were quantified with a SEAL QuAAtro analyzer (Bran + Luebbe, Germany; nutrients) and an Apollo SciTech DIC analyzer (Apollo SciTech, USA; DIC) as described in (Stratmann et al., 2026). The δ^13^C composition of DIC was determined with a Thermo Delta V continuous flow IRMS as reported in (Stratmann et al., 2026).

### Data analysis

#### Inorganic nutrient and dissolved inorganic carbon flux calculations

Benthic community fluxes of inorganic nutrients (nitrate, nitrite, ammonia, dissolved silicate, phosphate) [*Flux*_*benth com (i.nut)*_ (mmol inorg. nutrient m^-2^ d^-1^)] and DIC [*Flux*_*benth com (DIC)*_ (mmol DIC m^-2^ d^-1^)] were calculated as follows:

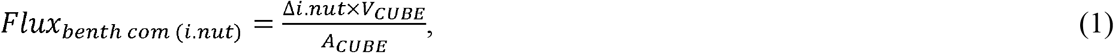

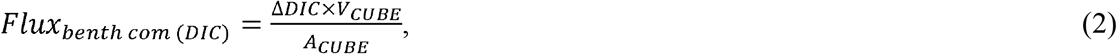

where Δ*i.nut* (μmol l^-1^ d^-1^) and Δ*DIC* (μmol l^-1^ d^-1^) are the changes in inorganic nutrient and DIC concentrations over time (T_0_ to T_6_) estimated via robust regression analyses (Western, 1995) (α = 0.05) using the ‘MASS’ package (Venables and Ripley, 2002). *V*_*CUBE*_ is the volume of the CUBE (=125 l) and *A*_*CUBE*_ is the surface area covered by the CUBE (=0.25 m^2^).

The benthic community flux of C-DIC 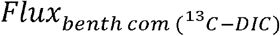 (mmol ^13^C-DIC m^-2^ d^-1^) was calculated as:

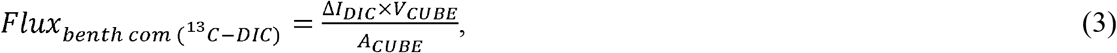

where Δ*I*_*DIC*_ (μmol ^13^C-DIC l^-1^ d^-1^) is the change of the production of ^13^C-DIC (*I*_*DIC*_) over time (T_0_ to T_6_) determined via robust regression analyses (α = 0.05).

#### Calculations of bulk ^13^C, ^15^N, and ^2^H uptake by macrozoobenthos

The bulk uptake 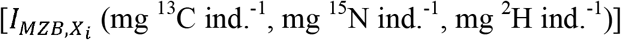 and biomass-specific uptake 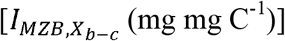 of ^13^C, ^15^N, and ^2^H by macrozoobenthos (MZB) were calculated as mentioned in (Stratmann et al., 2026). It was not necessary to correct for the exchange of noncovalently bound H in specimens from the experiments, because these specimens exchanged all noncovalently bound D back to H during the 14 d ‘turnover period’.

Total ^13^C, ^15^N, and ^2^H uptake of MZB [*TI*_*MZB,X*_ (mg m^-2^)] was calculated as:

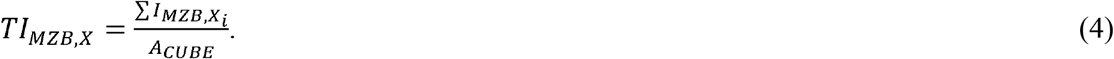

The ratios of ^13^C uptake/ ^15^N uptake 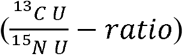, ^13^C uptake/ ^2^H uptake 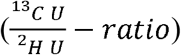, and ^15^N uptake/ ^2^H uptake 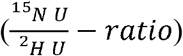 were calculated as:

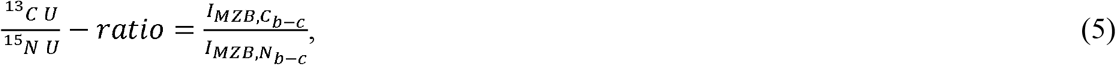

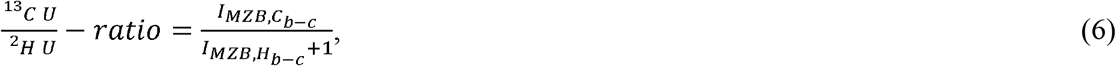

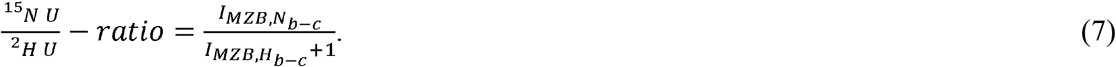

Changes in bulk ^13^C, ^15^N, and H uptake in MZB over time per season 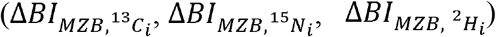 were calculated as:

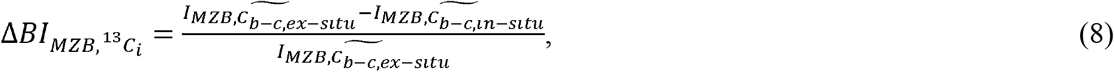

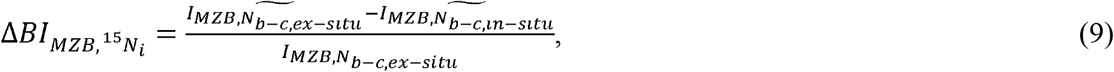

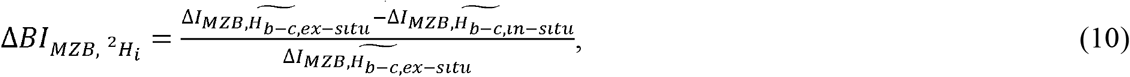

Where 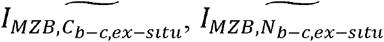, and 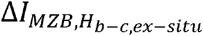 are the median biomass-specific bulk ^13^C, ^15^N, and ^2^H uptakes of a specific macrozoobenthos species and season (summer vs. autumn 2020) from (Stratmann et al., 2026). 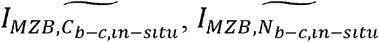, and 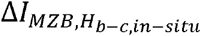 are the median biomass-specific bulk ^13^C, ^15^N, and ^2^H uptakes of a specific macrozoobenthos species and season from this study.

#### Calculation of ^13^C and ^2^H incorporation into sedimentary PLFAs

The incorporation of ^13^C (*I*_*C-PLFA,sed*_) and ^2^H (*I*_*H-PLFA,sed*_) into sedimentary PLFAs (µg ^13^C g^-1^ dry mass DM sediment; µg ^2^H g^-1^ DM sediment) was calculated as follows:

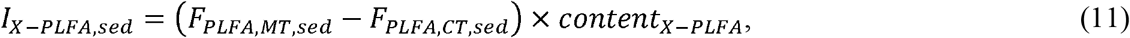

where *X* is the element C or H, *F*_*PLFA,MT,sed*_ is the fraction of the heavy stable isotope in sedimentary PLFAs from the main treatment, *F*_*PLFA,CT,sed*_ is the fraction of the heavy stable isotope in sedimentary PLFAs from the control, and *content*_*X-PLFA*_is the C and H content in sedimentary PLFAs.

#### Calculation of metabolic turnover rate of macrozoobenthos

Metabolic turnover rate *MTR* (% d^-1^) was calculated as:

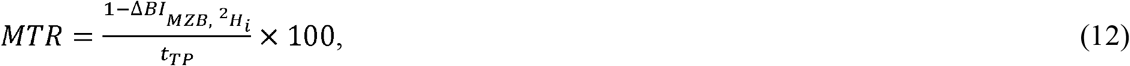

where *t*_*TP*_ is the ‘turnover period’ of the experiment (i.e., 14 d).

#### Statistical data analysis

Differences in *Flux*_*benth com*_ of inorganic nutrients, DIC, and ^13^C-DIC among sampling season (summer 2020, autumn 2020) and incubation time (day incubation, night incubation) were tested as changes in concentration over time by robust regression analysis. A flux was considered significant for a specific treatment (summer 2020, night incubation, *n* = 3; summer 2020, day incubation, *n* = 3; autumn 2020, night incubation, *n* = 6), when the *F*-statistic (α = 0.05) of the treatment was significant.

Differences in total ^13^C, ^15^N, and ^2^H uptake (*TI*_*MZB,C*_, *TI*_*MZB,N*_, *TI*_*MZB,H*_) among sampling seasons (summer 2020, autumn 2020) were tested with Welch’s t-tests (*t*; α = 0.05) in case of normally distributed data and with Mann-Whitney-Wilcoxon test (*V*, α = 0.05) for non-normally distributed data, after normality was assessed by the Shapiro-Wilk normality test.

Differences in biomass-specific ^2^H uptake among life stages (juvenile/ small vs. adult) of the same taxon (minimum *n*_juvenile_ = 3, *n*_adult_ = 3) in the experiments were tested with Mann-Whitney-Wilcoxon tests (*W*, α = 0.05) after Shapiro-Wilk normality tests showed that the data were not-normally distributed.

Differences in 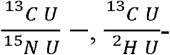, and 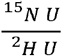-ratios among different life stages (juvenile/ small vs. adult) of the same taxon (minimum *n*_juvenile_ = 3, *n*_adult_ = 3) in the experiments were tested with Welch’s t-tests (*t*; α = 0.05; normally distributed data according to the Shapiro-Wilk normality test) and Mann-Whitney-Wilcoxon test (*W*, α = 0.05; not-normally distributed data according to the Shapiro-Wilk normality test), respectively. Data are presented as mean ± SD (median).

#### Fluxes of inorganic nutrients and dissolved inorganic carbon

Benthic community fluxes *Flux*_*benth com*_ of N-containing inorganic nutrients (i.e., nitrate, nitrite, and ammonia) varied substantially among sampling season and incubation period (Supplementary Figure 5), but most fluxes did not differ significantly from 0 mmol m^-2^ d^-1^ (Supplementary Table 5). Nitrate and nitrite were consumed during the day, but released during the night. Ammonia was consumed during the day, but released during the night in summer 2020, whereas phosphate was released during the night in summer, but consumed during the night in autumn. DIC was always produced by the benthic community, but ^13^C-DIC was taken up during the day and night in summer, while it was released in autumn.

#### Carbon and hydrogen incorporation into sedimentary PLFAs

Phospholipid-derived fatty acids known from cyanobacteria (i.e., C18:1ω9*c*) contained 5.13 10^-2^ ± 3.12 10^-2^ (4.45 10^-2^) ng ^13^C g DM sediment^-1^ and 5.01 10^-3^ ± 5.34 10^-3^ (2.76 10^-3^) ng ^2^H g DM sediment^-1^ in the upper 3 cm of sediment in summer 2020 and 2.54 10^-2^ ± 2.80 10^-2^ (1.94 10^-2^) ng ^13^C g DM sediment^-1^ and 5.13 10^-5^ ± 3.12 10^-5^ (4.45 10^-5^) ng ^2^H g DM sediment^-1^ in autumn 2020 (Fig. 2, 3; Supplementary Table 7). In the lower sediment layer (3 – max. 8 cm sediment depth), they contained 3.32 10^-7^ ± 8.12 10^-7^ (0.00) ng ^13^C g DM sediment^-1^ in summer and 5.20 10^-7^ ± 1.27 10^-6^ (0.00) ng ^13^C g DM sediment^-1^ in autumn 2020, but no ^2^H (Fig. 2, 3; Supplementary Table 7).

**Figure 2.**
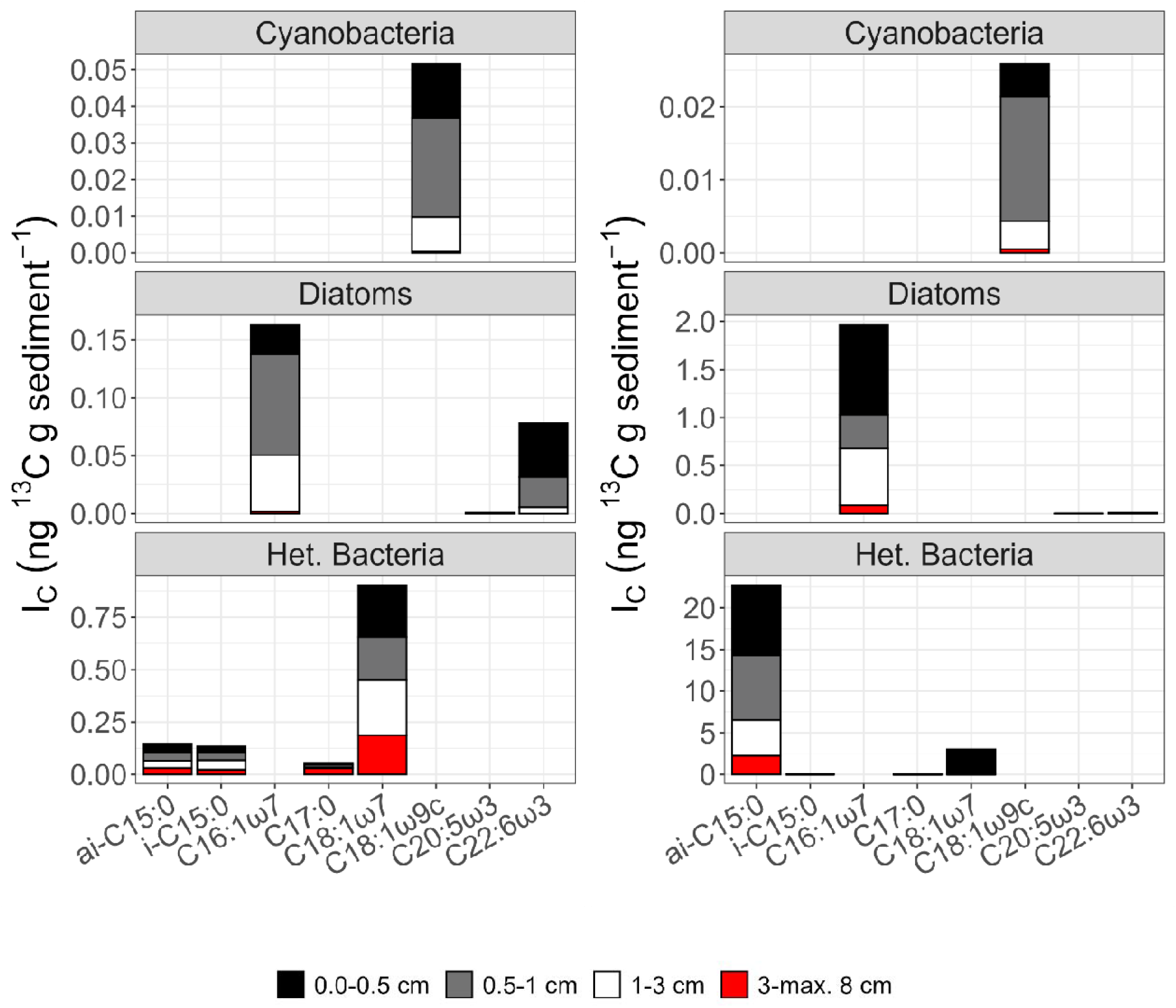
Incorporation of ^13^C (*I*_*C*_) into phospholipid-derived fatty acids (PLFA; ng ^13^C g DM sediment^-1^) typical for benthic cyanobacteria (C18:1ω9*c*), diatoms (C16:1ω7, C20:5ω3, C22:6ω3), and heterotrophic bacteria (*i*-C15:0 co-eluted with C14:1ω5*c*, C17:0, C18:1ω7) in summer (left panels) and autumn 2020 (right panels).

**Figure 3.**
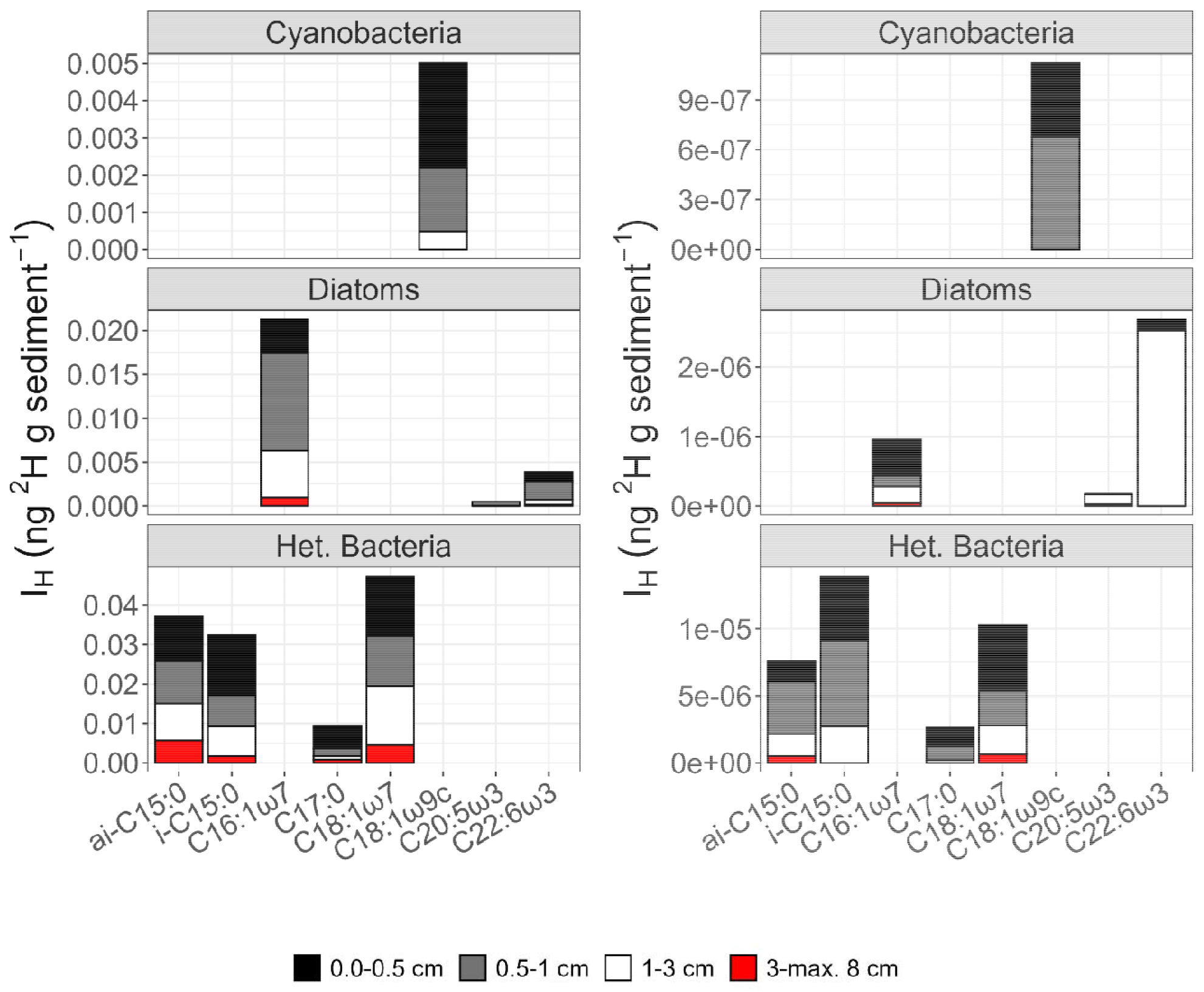
Incorporation of ^2^H (*I*_*H*_) into phospholipid-derived fatty acids (PLFA; ng ^2^H g DM sediment^-1^) typical for benthic cyanobacteria (C18:1ω9*c*), diatoms (C16:1ω7, C20:5ω3, C22:6ω3), and heterotrophic bacteria (*i*-C15:0 co-eluted with C14:1ω5*c*, C17:0, C18:1ω7) in summer (left panels) and autumn 2020 (right panels).

The amount of ^13^C and ^2^H that was incorporated into C18:1ω9*c* differed significantly between sediment layers (0 – 3 cm vs 3 – max. 8 cm) in summer 2020 (^13^C-incorporation: *χ*^*2*^(3) = 12.6, *p*-value = 0.006; *Z* = -3.39, *p*-value = 0.002); ^2^H-incorporation: *χ*^*2*^(3) = 10.8, *p*-value = 0.013; *Z* = - 3.51, *p*-value = 0.001) (Supplementary Table 8). There was also a significant difference between the upper sediment layer (0 – 3 cm) in summer 2020 and the lower sediment layer (3 – max. 8 cm) in autumn 2020 (^13^C-incorporation: *Z* = -3.35, *p*-value = 0.002; ^2^H-incorporation: *Z* = - 3.51, *p*-value = 0.001) (Supplementary Table 8).

Diatom-related PLFAs (C16:1ω7, C20:5ω3, C22:6ω3) in the upper 3 cm of sediment included 0.24 ± 0.25 (0.16) ng ^13^C g DM sediment^-1^ and 2.45 10^-2^ ± 3.44 10^-2^ (7.65 10^-3^) ng ^2^H g DM sediment^-1^ in summer 2020 and 1.89 ± 1.56 (1.89) ng ^13^C g DM sediment^-1^ and 3.75 10^-3^ ± 6.57 10^-3^ (1.17 10^-3^) ng ^2^H g DM sediment^-1^ in autumn 2020 (Fig. 2, 3; Supplementary Table 7). In the 3 – max 8 cm sediment layer, these PLFAs still contained 1.84x10^-3^ ± 2.38x10^-3^ (9.71x10^-4^) ng ^13^C g DM sediment^-1^ and 1.13x10^-3^ ± 2.09x10^-3^ (2.25x10^-4^) ng ^2^H g DM sediment^-1^ in summer 2020 and 5.20x10^-4^ ± 1.27x10^-3^ (0.00) ng ^13^C g DM sediment^-1^ and 6.56x10^-5^ ± 9.62x10^-5^ (1.16x10^-3^) ng ^2^H g DM sediment^-1^ in autumn 2020 (Fig. 2, 3; Supplementary Table 7). Diatom-related PLFAs included significantly different concentrations of ^13^C in the two sediment layers (0 – 3 cm vs 3 – max. 8 cm) in autumn 2020 (*χ*^*2*^(3) = 12.8, *p*-value = 0.005; *Z* = - 3.06, *p*-value = 0.007), and the incorporated ^13^C concentrations differed significantly between the lower sediment layer (3 – max. 8 cm) in summer vs. the upper sediment layer (0 – 3 cm) in autumn 2020 (*Z* = 2.77, *p*-value = 0.02) (Supplementary Table 8). The concentration of ^2^H in PLFA from diatoms only differed between upper sediment layer (0 – 3 cm) in summer 2020 and lower sediment layer (3 – max. 8 cm) in autumn 2020 (*χ*^*2*^(3) = 11.5, *p*-value = 0.009; *Z* = -3.57, *p*-value = 0.36) (Supplementary Table 8).

PLFAs specific for sedimentary heterotrophic bacteria (*i*-C15:0 co-eluted with C14:1ω5*c*, C17:0, C18:1ω7) consisted of 0.97 ± 0.53 (0.89) ng ^13^C g DM sediment^-1^ (0 – 3 cm) and 0.27 ± 0.18 (0.22) ng ^13^C g DM sediment^-1^ (3 – max. 8 cm) in summer 2020 and of 23.6 ± 17.1 (23.9) ng ^13^C g DM sediment^-1^ (0 – 3 cm) and 2.24 ± 2.52 (1.95) ng ^13^C g DM sediment^-1^ (3 – max. 8 cm) in autumn 2020 (Figure 2, Supplementary Table 8). The differences in ^13^C incorporated into these heterotrophic bacteria-specific PLFA was not significant, neither between seasons nor sediment layers (*χ*^*2*^(3) = 7.45, *p*-value = 0.06). Additionally, these PLFAs included 0.11 ± 0.05 (0.11) ng ^2^H g DM sediment^-1^ (0 – 3 cm) and 1.26x10^-2^ ± 8.83x10^-3^ (1.35x10^-2^) ng ^2^H g DM sediment^-1^ (3 – max. 8 cm) in summer 2020 and 3.32x10^-2^ ± 3.04x10^-3^ (3.67x10^-2^) ng ^2^H g DM sediment^-1^ (0 – 3 cm) and 1.13x10^-3^ ± 1.59x10^-3^ (2.28x10^-4^) ng ^2^H g DM sediment^-1^ (3 – max. 8 cm) in autumn 2020 (Figure 3, Supplementary Table 8). The difference in ^2^H concentrations present in the heterotrophic bacteria-specific PLFAs was only significant for the 3 – max. 8 cm sediment layer in autumn vs. the 0 – 3 cm sediment layer in summer 2020 (*χ*^*2*^(3) = 14.2, *p*-value = 0.003; *Z* = -3.99, *p*-value = 0.0002) (Supplementary Table 8).

### Total macrozoobenthos uptake of carbon, nitrogen, and hydrogen

Macrozoobenthos took in total 87.4 ± 86.5 (54.5) mg ^13^C m^-2^, 47.0 ± 42.6 (34.0) mg ^15^N m^-2^, and 7.69x10^-2^ ± 7.13x10^-2^ (4.98x10^-2^) mg ^2^H m^-2^ up in summer 2020 and 60.0 ± 48.3 (54.9) mg ^13^C m^-2^, 26.8 ± 20.8 (24.2) mg ^15^N m^-2^, and 8.42x10^-2^ ± 6.51x10^-2^ (6.04x10^-2^) mg ^2^H m^-2^ in autumn 2020. The difference in uptake of ^13^C, ^15^N, and ^2^H between the two seasons, however, was not significant (^13^C: *t*(7.84) = 0.68, *p*-value = 0.52; ^15^N: *t*(7.24) = 1.05, *p*-value = 0.33; ^2^H: *t*(9.92) = -0.18, *p*-value = 0.86).

In summer 2020 most ^13^C was taken up by *H. panicea, M. gigas*, and *C. fornicata* (97.9% of total ^13^C-uptake), whereas most ^15^N and ^2^H was incorporated into *H. panicea, M. gigas*, and *R. philippinarum* (98.1% of total ^15^N-uptake, 92.8% of total ^2^H-uptake) (Supplementary Figure 6; Supplementary Table 9). In comparison, in autumn 2020 *H. panicea, H. perlevis*, and *M. gigas* dominated ^13^C-uptake (93.3% of total ^13^C-uptake) and ^15^N-uptake (93.3% of total ^15^N-uptake), and ^2^H was incorporated into tissue of predominantly *H. perlevis, H. panicea*, and *M. gigas* (87.7% of total ^2^H-uptake).

#### Biomass-specific uptake of carbon, nitrogen, and hydrogen by the benthic community

Biomass-specific ^13^C uptake in summer 2020 was dominated by *H. panicea, C. fornicata, R. philippinarum, Malacoceros fuliginosus* (64.4% of total ^13^C-uptake), and by *H. panicea, H. perlevis, C. fornicata*, and unidentified Polychaeta in autumn (58.5% of total ^13^C-uptake) (Fig. 4). Most ^15^N was incorporated by *H. panicea, R. philippinarum, C. fornicata*, and *M. fuliginosus* in summer (62.8% of total ^15^N-uptake), and by *H. panicea, H. perlevis, C. fornicata*, and unidentified Polychaeta in autumn (58.3% of total ^15^N-uptake), while *Rhithropanopeus harrisii, L. conchilega*, and *Owenia fusiformis* incorporated most ^2^H in summer (81.4% of total ^2^H-uptake) and *R. harrisii, H. perlevis, C. fornicata, E. longa, H. panicea*, and *Psammechinus miliaris* consumed most ^2^H in autumn (46.4% of total ^2^H-uptake) (Fig. 4).

**Figure 4.**
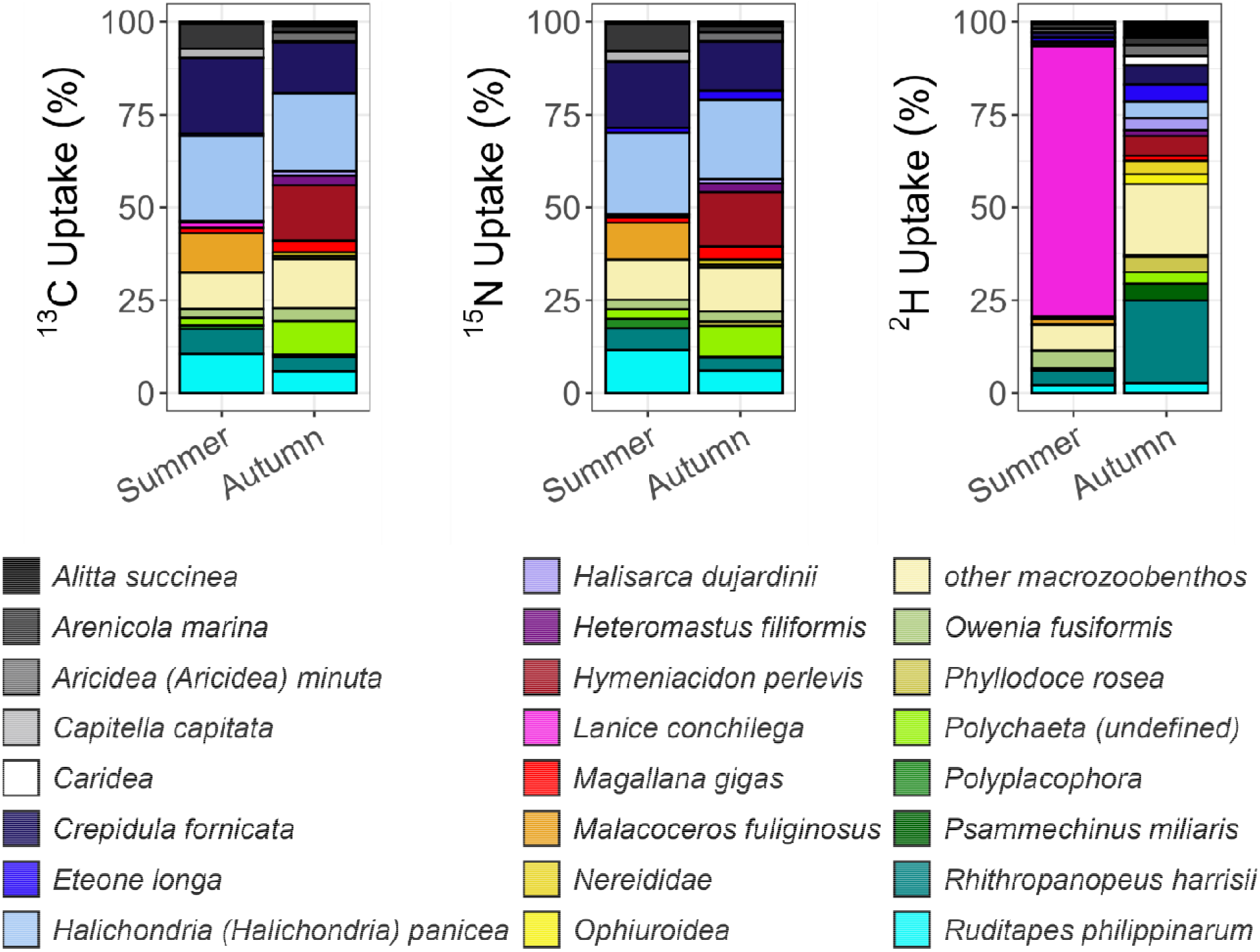
Relative biomass-specific uptake of ^13^C, ^15^N, and ^2^H (% total biomass-specific uptake) by macrozoobenthos in summer and autumn 2020. Macrozoobenthos taxa that took up ≤2.5% of total biomass-specific ^13^C, ^15^N, and/ or ^2^H uptakes are aggregated under the category “other macrozoobenthos”.

When separated by life stage (adult vs. juveniles/ small specimens), the lowest biomass-specific ^2^H uptake rates were measured in adult *Eumida sanguinea* (3.08 ± 2.71 (4.12) ng ^2^H mg C^-1^) and in juvenile/small *Nephtys cirrosa* (0.25 ng ^2^H mg C^-1^). In comparison, the highest rates were detected in *E. longa* (20.6 ± 21.0 (12.6) ng ^2^H mg C^-1^; adult stage) and in *O. fusiformis* (58.2 ± 80.2 (58.2) ng ^2^H mg C^-1^; juvenile/ small stage) (Supplementary Table 10). Additionally, a direct comparison between juvenile/ small and adult life stages of taxa (minimum *n* = 3 per life stage) revealed that only the biomass-specific ^2^H uptake by *R. harrisii* juveniles differed from *R. harrisii* adults (*W* = 144, *p* = 0.049). The life stage-dependent ^2^H uptake rates of all other investigated taxa were not significantly different (Supplementary Table 11).

#### ^13^C/^15^N, ^13^C/^2^H, and ^15^N/^2^H ratios

The 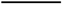-ratios of macrozoobenthos ranged from 0.17 ± 0.38 (0.00) (Anthozoa) to 204 ± 934 (1.66) (*C. maenas*) in summer 2020 (Supplementary Figure 7) and from 0.15 ± 0.25 (0.00) (*Palaemon adspersus*) to 27.0 (Polyplacophora) in autumn 2020 (Supplementary Figure 7). In summer 2020, the 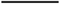 - and 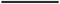 -ratios of macrozoobenthos were highest for *M. fuliginosus* (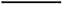 -ratio: 2.17 10^-3^, 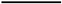 -ratio: 1.09 10^-3^) and lowest for Anthozoa (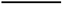 -ratio: 9.00 10^-6^) and Ophiuroidea (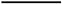 -ratio: 1.00 10^-6^) (Supplementary Figure 7). In comparison, in autumn 2020, the 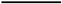- and 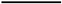 -ratios of macrozoobenthos ranged from *P. adspersus* (3.00 10^-6^ ± 5.00 10^-6^ (0.00)) and Amphipoda (1.00 10^-6^), respectively to *H. panicea* (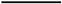 -ratio: 1.40 10^-3^ ± 9.04 10^-4^ (1.33 10^-3^), 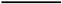 -ratio: 6.32 10^-4^ ± 3.88 10^-4^ (6.38 10^-4^)) (Supplementary Figure 7).

When feeding types were considered, the median 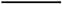 -ratios increased from carnivores, scavengers, deposit feeders, herbivores, omnivores, to filter- and suspension feeders (Fig. 5). The feeding types with the highest median 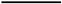 - and 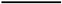 -ratios were filter and suspension feeders, followed by deposit feeders and carnivores (Fig. 5).

**Figure 5.**
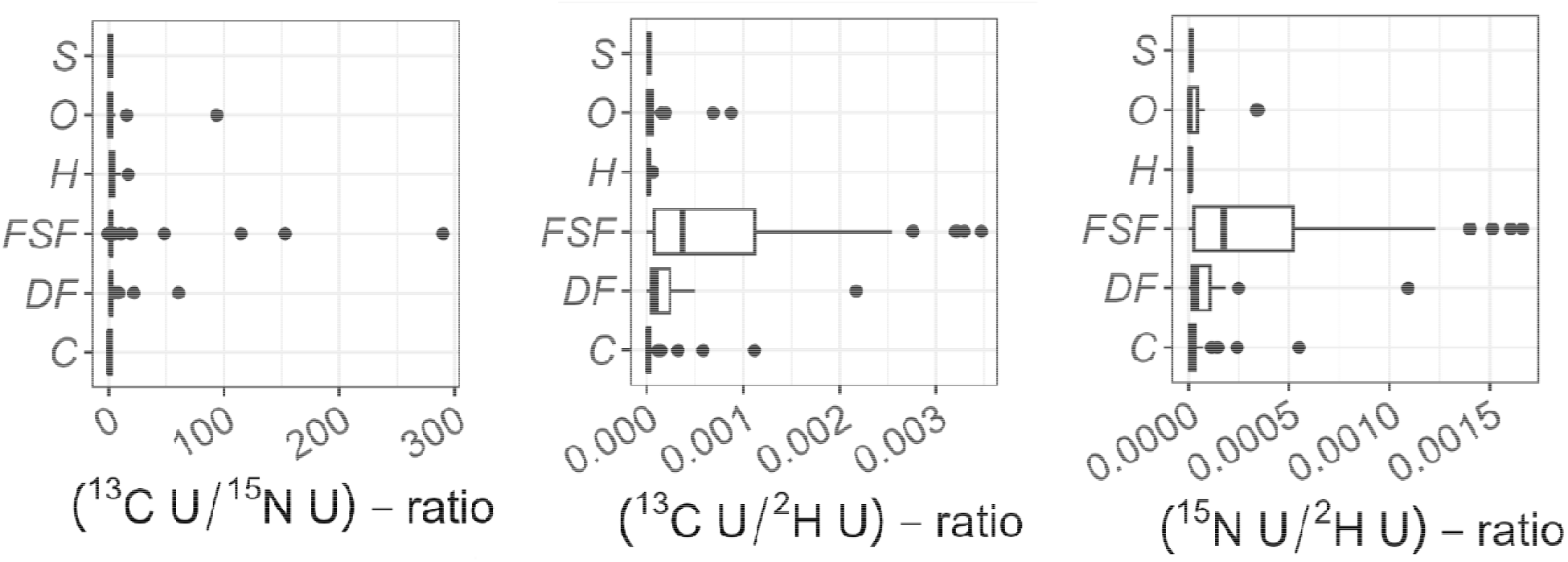
Ratios of 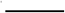, 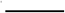, and 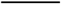 of adult macrozoobenthos belonging to different feeding types. Note that ratios were only shown for specimens that took up ^13^C, ^15^N, and/ or ^2^H and that were identified to species level. Species were classified in feeding types based on published literature (Supplementary Table 12). Abbreviations: C = carnivore, DF = deposit feeder, FSF = filter and suspension feeder, H = herbivore, O = omnivore, S = scavenger.

Assessing adult locomotion showed that the highest median 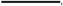 -ratio were detected in sessile species, followed by crawling/ creeping/ climbing species, burrowing and swimming species (Fig. 6). Median 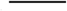 - and 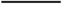 -ratios were highest for sessile, burrowing, swimming, and crawling/ creeping/ climbing species (Fig. 6).

**Figure 6.**
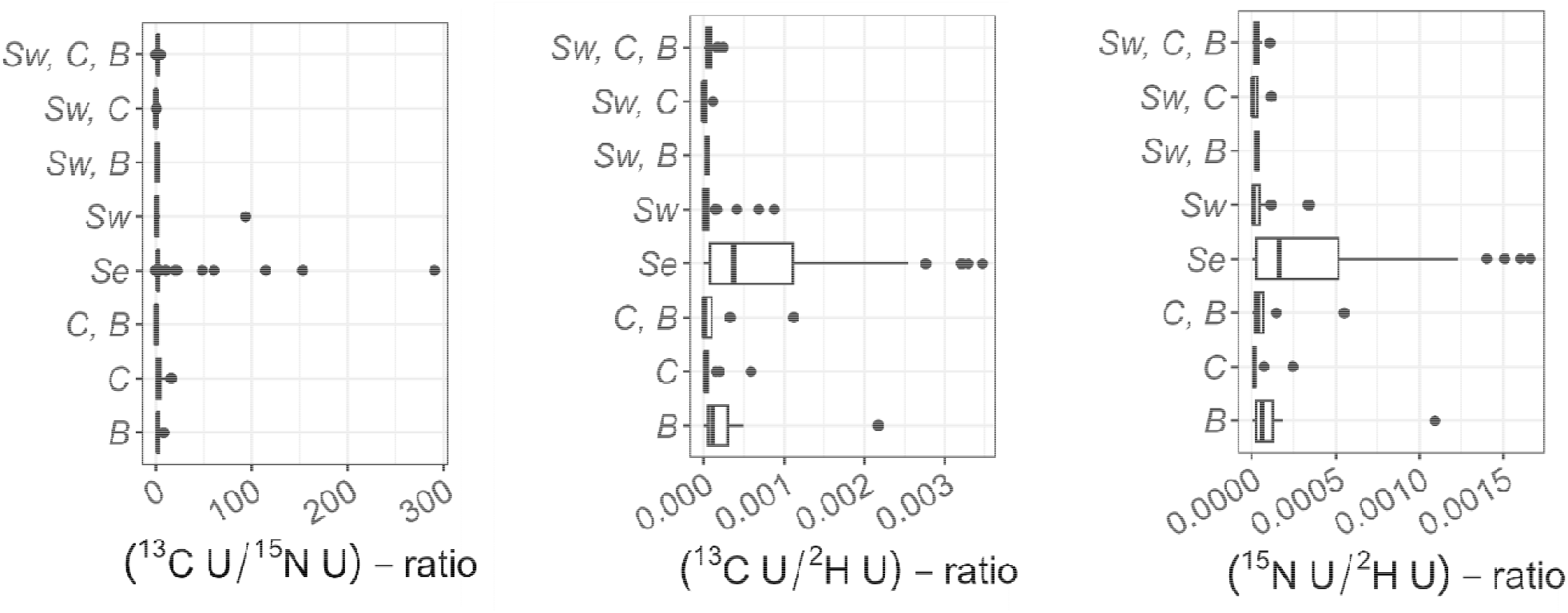
Ratios of 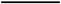, 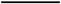, and 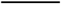 of adult macrozoobenthos having different motility types. Species were classified in motility types based on published literature (Supplementary Table 12). Abbreviations: B = burrower, C = crawl/ creep/ climb, Se = sessile, Sw = swimming.

A comparison of life stages of species (Fig. 7) for which minimum 3 replicates per life stage were available revealed that the 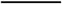 - and 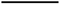 -ratios of juveniles and adults were only different for *R. harrisii* (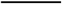 : *W* = 309, *p* = 0.003, 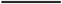 : *W* = 5, *p* < 0.001) (Supplementary Table 13). Significant differences between juveniles and adults of the same species existed for 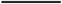 -ratios in case of *C. fornicata* (*W* = 35, *p* = 0.016), *E. longa* (*W* = 0, *p* = 0.008), and *R. harrisii* (*W* = 0, *p* < 0.001) (Supplementary Table 13).

**Figure 7.**
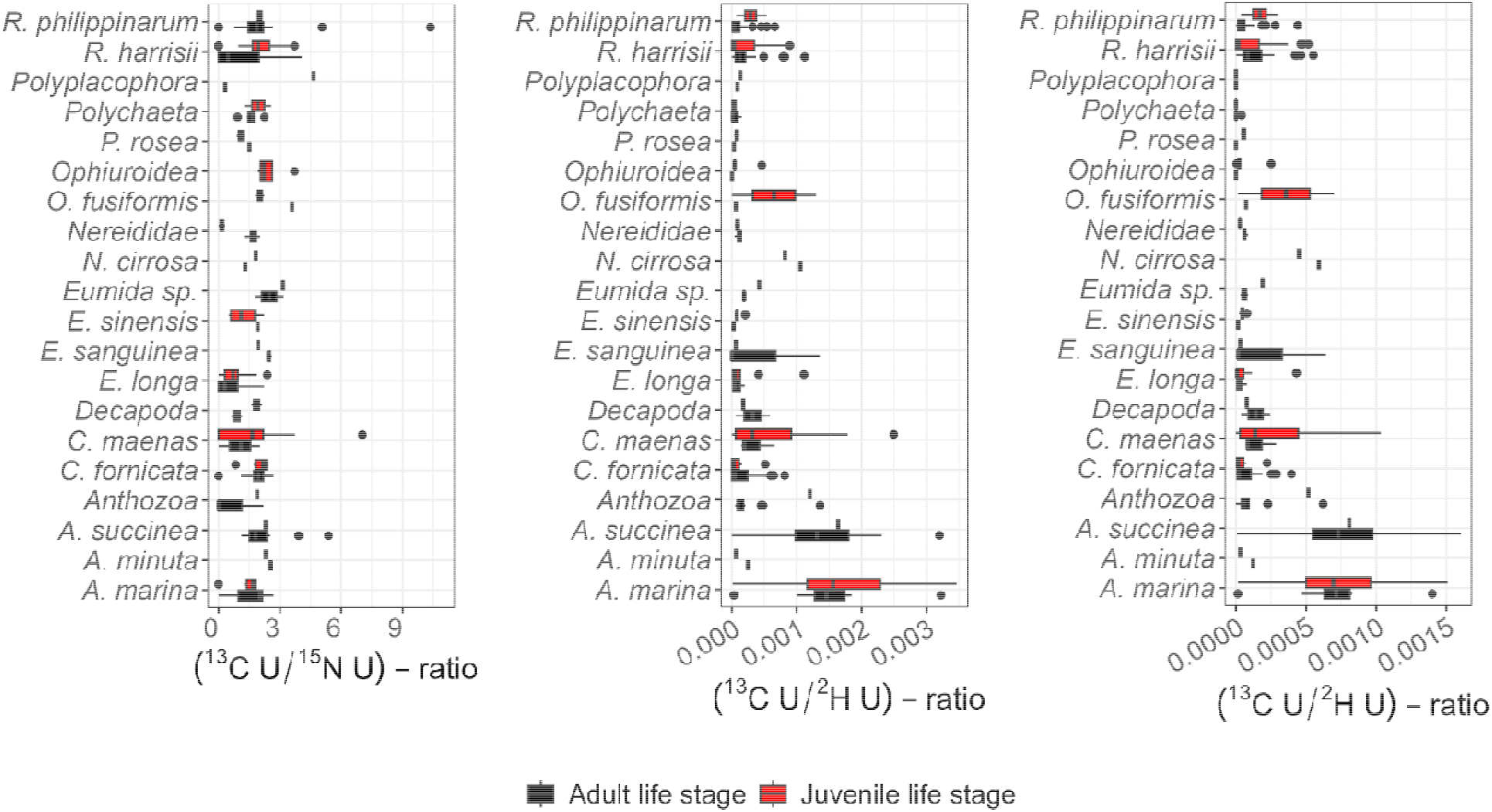
Ratios of 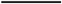, 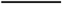, and 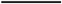 of juvenile and adult specimens of macrozoobenthos taxa, excluding very high data points in 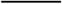 -ratio. Note that ratios were only shown for taxa that were present in juvenile and adult life stages.

#### Changes in bulk ^13^C, ^15^N, and ^2^H uptake over time

The difference in average bulk ^13^C uptake by macrozoobenthos between *ex-situ* (Stratmann et al., 2026) and *in-situ* experiments, the so-called change in bulk ^13^C uptake over time (Δ*BI*), ranged from -2.38 µg ^13^C mg C^-1^ (*H. panicea*) in summer and -0.25 µg ^13^C mg C^-1^ (*M. gigas*) in autumn 2020 to 1.00 µg ^13^C mg C^-1^ (*P. depurator*) in summer and 1.00 µg ^13^C mg C^-1^ (*P. depurator*) in autumn 2020 (Fig. 8, upper panel). The change in bulk ^15^N uptake over time was lowest in *H. panicea* (-2.64 µg ^15^N mg C^-1^, summer 2020) and *M. gigas* (-0.28 µg ^15^N mg C^-1^, autumn 2020) and highest in *E. sinensis* (0.86 µg ^15^N mg C^-1^, summer 2020) and *L. littorea* (0.96 µg ^15^N mg C^-1^, autumn 2020) (Fig. 8, middle panel), whereas the change in bulk ^2^H uptake over time ranged from 0.33 µg ^2^H mg C^-1^ (*H. panicea*, summer 2020) and 0.47 µg ^2^H mg C^-1^ (*H. panicea*, autumn 2020) to 1.00 µg ^2^H mg C^-1^ (*P. depurator*, summer 2020) and 1.00 µg ^2^H mg C^-1^ (*L. littorea*, autumn 2020) (Fig. 8, lower panel).

**Figure 8.**
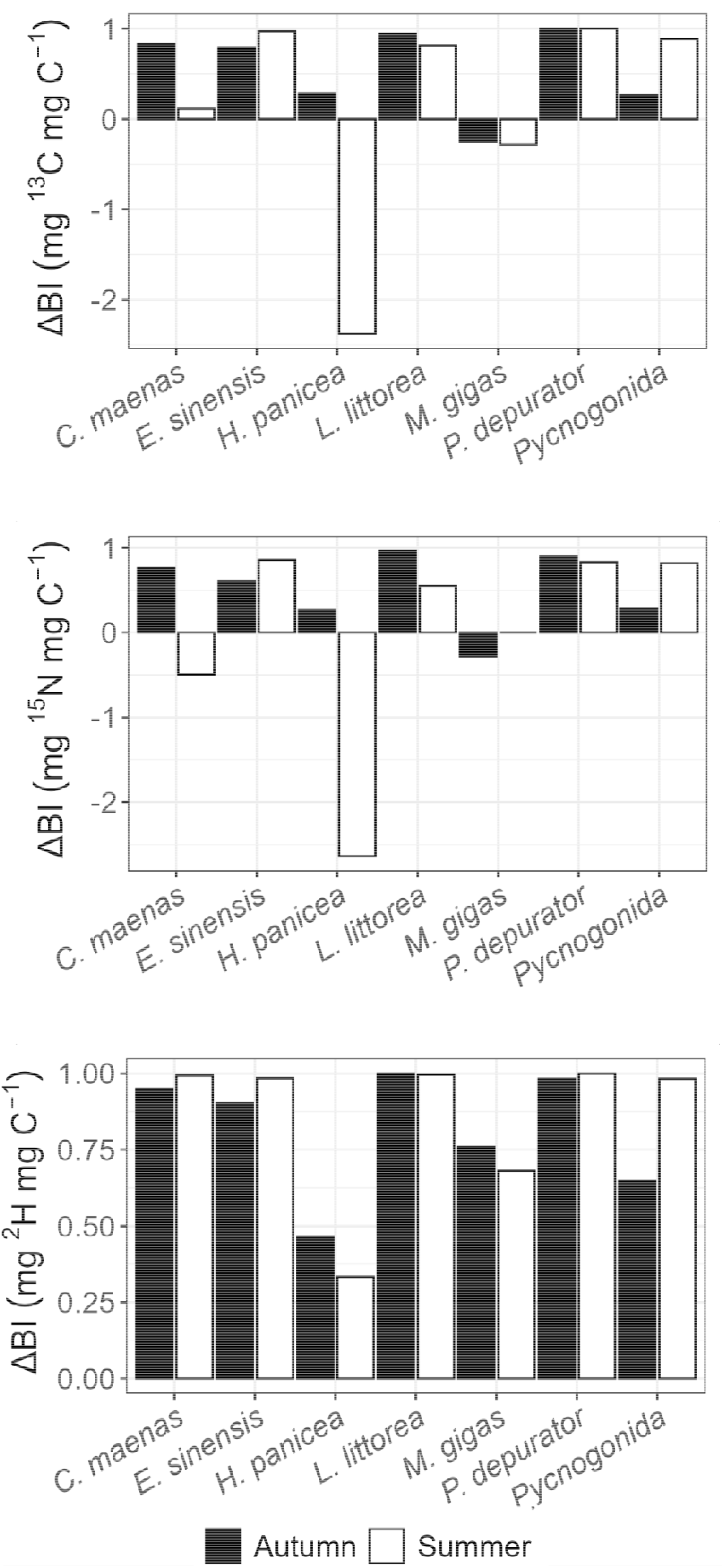
Changes in bulk ^13^C, ^15^N, and ^2^H uptake by macrozoobenthos over time (, mg ^13^C mg C^-1^;, mg ^15^N mg C^-1^;, mg ^2^H mg C^-1^) in summer and autumn 2020 relative to the short-term *ex-situ* experiment (Stratmann et al., 2026).

#### Metabolic turnover rate

The *MTR* ranged from 0.04 % d^-1^ in summer (*L. littorea*) and 0.13 % d^-1^ autumn 2020 (*P. depurator*) to 4.76 % d^-1^ (*H. panicea*) in summer and 3.82 % d^-1^ (*H. panicea*) in autumn 2020 (Fig. 9). *MTR* for *P. depurator* in summer 2020 and *L. littorea* in autumn 2020 could not be determined, because no ^2^H uptake was detected.

**Figure 9.**
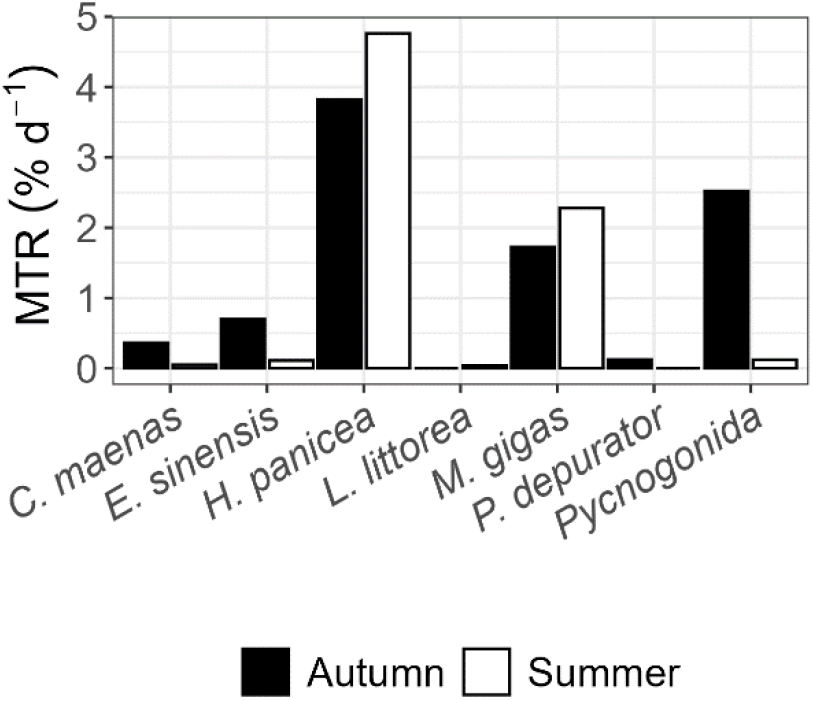
Metabolic turnover rate *MTR* (% d^-1^) of various macrozoobenthos species present in summer and autumn 2020.

## Discussion

### Long-term benthic community responses to pelagic bacteria

Fourteen days after feeding the intertidal and Pacific oyster community with ^13^C and ^15^N-enriched substrate bacteria, most ^13^C and ^15^N per biomass was detected in sessile filter and suspension feeders (i.e., *H. panicea, C. fornicata, H. perlevis, R. philippinarum*) (Fig. 10).

**Figure 10.**
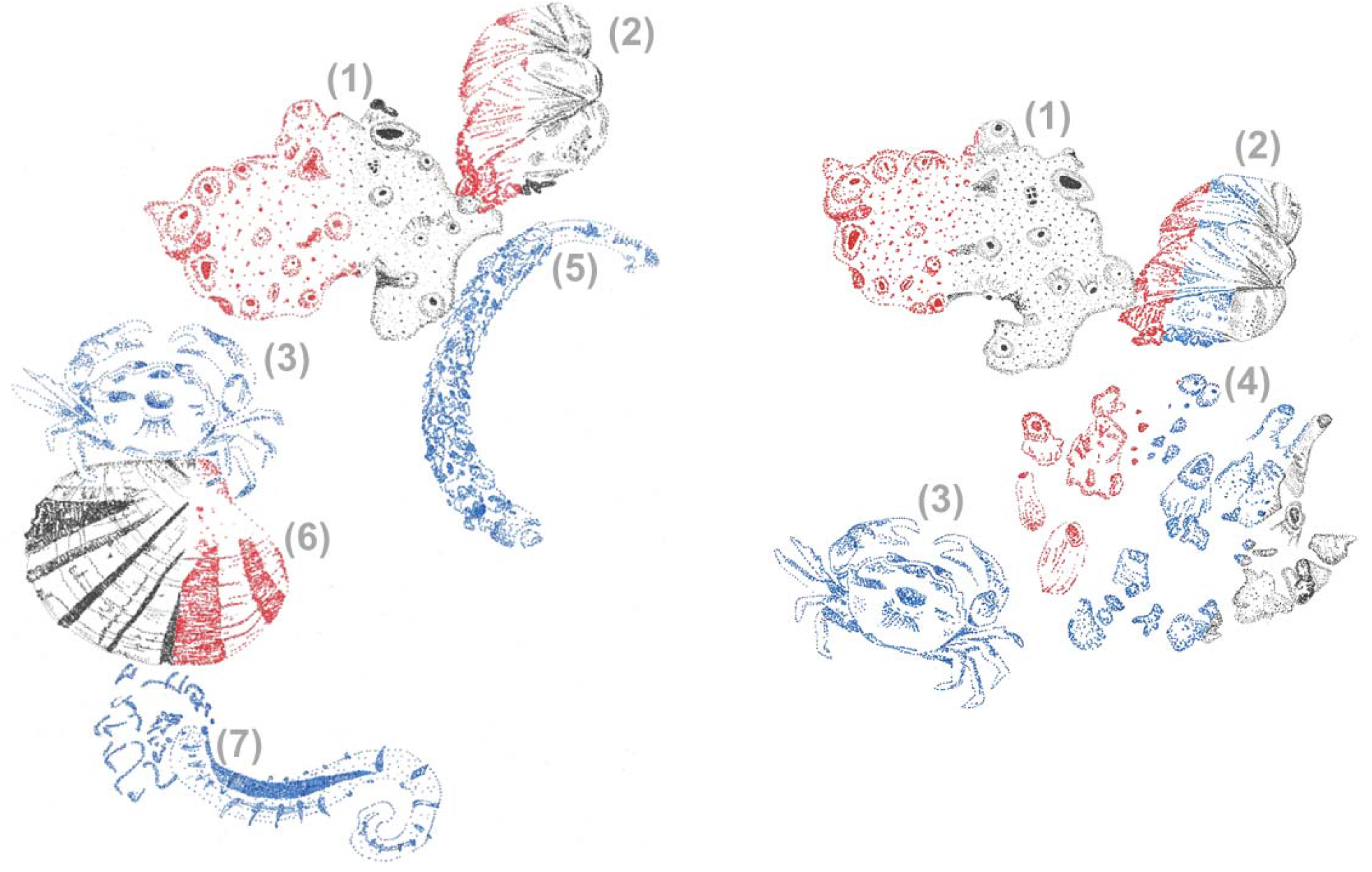
Drawings of the three macrozoobenthos species that took up most ^15^N (red), ^2^H (blue), and ^13^C (black) on a biomass (mg C) level per season (left panel) summer vs. (right panel) autumn 2020. A multicolor drawing of a species indicates that this species belongs the three species with the highest uptake for multiple stable isotopes. The species presented are: (1) *Halichondria* (*Halichondria*) *panicea*, (2) *Crepidula fornicata*, (3) *Rhithropanopeus harrisii*, (4) *Hymeniacidon perlevis*, (5) *Owenia fusiformis*, (6) *Ruditapes philippinarum*, and (7) *Lanice conchilega*. Illustrations by Tanja Stratmann.

***Halichondria panicea*** is the sponge species that took up most ^13^C and ^15^N-enriched substrate bacteria in the short-time (<12 h) *ex-situ* incubation experiments (Stratmann et al., 2026). Even after 14 d, ^13^C and ^15^N derived from substrate bacteria was still measurable in *H. panicea* in the *in-situ* experiments (Supplementary Table 14). However, this species may process the ingested ^13^C and ^15^N differently. A comparison of the 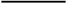 -ratio of *H. panicea* from the *ex-situ* (Stratmann et al., 2026) vs. *in-situ* experiments (this study) with the 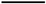 -ratio of substrate bacteria from the *ex-situ* vs. *in-situ* experiments indicates that *H. panicea* retained more substrate bacteria-derived N relative to C. Hence, the sponge respired more of the assimilated C as (^13^C)-DIC than it used for the production of sponge (holobiont) biomass, whereas it retained more of the assimilated N for biomass production than it mineralized/ ammonificated to (^15^N)-NH_4_^+^. This preferential retainment of N over C did not change over a 14 d period and has been observed in other sponge species: For instance, the massive demosponge *Spongosorites coralliophaga*, present in cold-water coral reefs, preferentially assimilated and retained N over C when fed with ^13^C- and ^15^N-enriched microalgae and bacteria in a dual stable isotope tracer study (Kazanidis et al., 2018). Other deep-sea sponges (i.e., *Geodia barretti, Hymedesmia* (*Hymedesmia*) *paupertas, Vazella pourtalesii*) assimilate more N from ^13^C- and ^15^N-enriched bacteria than from ^13^C- and ^15^N-enriched DOM (Bart et al., 2020). As a result, (Bart et al., 2020) speculated that this might be a strategy to maintain the stoichiometric homeostasis of the sponges. Unfortunately, it has not been tested whether *H. panicea* is able to assimilate DOM and therefore we do not know whether this intertidal/ subtidal species retains N differentially when it ingests DOM compared to when it ingests bacteria.

***Crepidula fornicata*** is a limpet that was present at the *in-situ* experimental sites in summer and autumn 2020, but in the *ex-situ* experiments it was only part of the macrozoobenthos epifauna in autumn (Stratmann et al., 2026). After 14 d, it still contained between 3.33 µg ^13^C mg C^-1^ (autumn) and 4.24 µg ^13^C mg C^-1^ (summer) and between 1.43 µg ^15^N mg C^-1^ (autumn) and 1.97 µg ^15^N mg C^-1^ (summer) (Supplementary Table 14), and therefore 35% less ^13^C and 28% less ^15^N than the specimens in the *ex-situ* experiments (Stratmann et al., 2026) in autumn 2020. When we assume that *C. fornicata* incorporated the same amount of ^13^C and ^15^N derived from substrate bacteria in the *in-situ* (this study) and *ex-situ* (Stratmann et al., 2026) experiments in autumn, they mineralized about 30% of the ingested substrate bacteria within two weeks. Assuming further that *C. fornicata* had mineralized ^13^C and ^15^N at similar rates in summer 2020, the species incorporated at least 5.72 µg ^13^C mg C^-1^ and 2.52 µg ^15^N mg C^-1^ at the beginning of the *in-situ* experiments.

Most of the ^13^C and ^15^N likely originated from the ^13^C and ^15^N-enriched substrate bacteria, though a smaller portion might have come from (suspended) microphytobenthos that had incorporated recycled label. Values of δ^13^C and δ^15^N of *C. fornicata* specimens collected at Bourgneuf Bay and at Bay of Mont Saint Michel (both France) indicated that the natural diet of this species is mostly comprised of phytoplankton and benthic diatoms (Decottignies et al., 2007; Riera, 2007): Whereas juvenile *C. fornicata* (<2.8 cm shell length) and adult mobile males graze upon substrate and are suspension feeders (Chaparro et al., 2002; Navarro and Chaparro, 2002), implying that they could take microphytobenthos and phytoplankton up directly, adult sessile female *C. fornicata* are exclusively suspension feeders (Chaparro et al., 2002; Navarro and Chaparro, 2002), signifying that they would incorporate phytoplankton, suspended microphytobenthos, and free-living bacteria.

Based on our data, we propose that macrozoobenthos at the experimental sites respired parts of the ^13^C they had ingested in the form of ^13^C-enriched substrate bacteria as ^13^C-DIC (Supplementary Figure 5). This ^13^C-DIC might have been used by microphytobenthos for primary production as indicated by ^13^C-incorporation into sedimentary PLFAs typical for diatoms and benthic cyanobacteria (Fig. 2). If only a fraction of this ^13^C-enriched microphytobenthos was suspended by the stirring inside the CUBEs or by mobile, bioturbating macrozoobenthos species inside the chambers, juvenile *C. fornicata*, adult male and even sessile suspension feeding female *C. fornicata* had access to this food source and could have ingested it. In case all the *C. fornicata* adult specimens were male, they might have grazed directly upon ^13^C-enriched microphytobenthos. Unfortunately, we did not separate male from female *C. fornicata* adults and therefore cannot test this, as we only separated juvenile *C. fornicata* life stages from adult ones (Fig. 7).

The sponge ***Hymeniacidon perlevis*** had high ^13^C and ^15^N uptake rates in autumn 2020, because the species is able to increase its food uptake with increasing food availability (Maldonado et al., 2010). It is also more efficient in taking up substrate bacteria compared to other benthic filter feeders (Longo et al., 2016). In summer 2020, *H. perlevis*’ uptake comprised 0 µg ^13^C mg C^-1^ and 0 µg ^15^N mg C^-1^, because the species was neither present at the experimental sites nor at the reference sites (Supplementary Table 9).

The non-native clam ***Ruditapes philippinarum*** was first found as complete halves in Yerseke at the Eastern Scheldt in 2007 (Faasse and Ligthhart, 2008) and since then it has dispersed into the Wadden Sea (Reise et al., 2024). This species filters and ingests mostly (nano)phytoplankton (e.g., dinoflagellates like *Karlodinium veneficum* and chlorophytes) and resuspended microphytobenthos (e.g., diatoms), but also sedimentary organic matter (SOM) and particulate organic matter (POM) (Kasai et al., 2004; Dang et al., 2009; Komorita et al., 2014; Dias et al., 2019; Liu et al., 2022). Occasionally, *R. philippinarum* also filters free-living bacteria out of the water column, but with lower filtration rates than it uses to filter picocyanobacteria (Nakamura, 2001).

At first glance, our study seems to confirm that clams take up free-living bacteria, as this species was highly enriched in ^13^C- and ^15^N derived from concentrated substrate bacteria (Fig. 4, 10; Supplementary Table 14). However, the 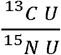-ratio of *R. philippinarum* was 2.13 ± 2.07 (1.77) in summer 2020 and 1.83 ± 1.10 (2.02) in autumn (Supplementary Table 14), and therefore 44% lower than the 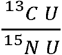 -ratio of the substrate bacteria (3.27 ± 0.34 (3.24)) in autumn (Supplementary Table 2). A lower 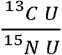 -ratio might suggest that *R. philippinarum* retained preferentially ^15^N because the species was N limited, but the 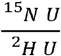 -ratio (Fig. 7) indicated otherwise. (Stratmann et al., 2026) showed that a high feeding activity combined with a median metabolic activity may lead to 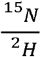 -ratio >1 which signifies a likely N limitation. In comparison, a reduced feeding activity combined with a high metabolic activity (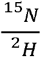 -ratio <1) implies a reduced N limitation (Stratmann et al., 2026). In this study, the 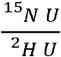 -ratio of *R. philippinarum* in autumn 2020 was 0.16 ± 0.18 (0.06) suggesting a low to no N limitation.

As the clam is known to ingest (re)suspended microphytobenthos (Dang et al., 2009; Dias et al., 2019), *R. philippinarum* might have fed upon suspended microphytobenthos. In fact, several PLFAs that are typical for cyanobacteria and diatoms (i.e., C16:1ω7, C18:1ω9c, C22:6ω3) were enriched in ^13^C (Fig. 2). This enrichment was likely the result of microphytobenthos taking up ^13^C-DIC derived from mineralized substrate bacteria. Unfortunately, we lack information about ^15^N enrichment of bulk microphytobenthos or about ^15^N enrichment of amino acids, so we cannot calculate the contribution of ^13^C and ^15^N enriched substrate bacteria vs. ^13^C and ^15^N enriched microphytobenthos, and provide more detailed information about the diet of *R. philippinarum* at the experimental and reference sites.

### Life stage differences in hydrogen processing

The largest differences in ^2^H uptake between juvenile/ small and adult specimens of the same species were detected in the crab ***R. harrisii*** (juvenile/ small life stage: 20.4 ± 27.7 ng ^2^H mg C^-1^; adult: 10.6 ± 20.9 ng ^2^H mg C^-1^; Supplementary Table 10). Young adult *R. harrisii* pass through four to five molt cycles in the first year with ∼10 d of intermold (Turoboyski, 1973). One to 4 h after molting, the body of *R. harrisii* grows (Turoboyski, 1973). This body growth may be linked to an increased ^2^H uptake. Hence, we propose that the two times higher ^2^H uptake by juvenile/ small R. *harrisii* compared to adult specimens was likely the result of post-molt growth.

Adult ***C. maenas***, in contrast, had a (not-significantly) higher ^2^H uptake rate compared to juvenile/ small specimens (juvenile/ small life stage: 3.33 ± 4.56 ng ^2^H mg C^-1^; adult: 5.38 ± 6.02 ng ^2^H mg C^-1^; Supplementary Table 10). Like *R. harrisii*, this species passes through five developmental stages prior to reaching the crab stage (Williams, 1967). After reaching this stage, *C. maenas* molds up to 18 times more during its lifetime (Crothers, 1967), but only once per year as soon as the crab reaches a size of ∼3 cm (Broekhuysen, 1937). *Carcinus maenas* can only grow until its carapace has hardened and the length of this post-molt growth period increases with increasing crab size from 3 – 5 d at 1.7 cm size to ∼5 d at 5.6 cm size (Broekhuysen, 1937). In 1935, the main molting period of the species was July, August, and September for females, and May and June for males in the Dutch Wadden Sea (Broekhuysen, 1937), a period of time that overlaps with the experimental period 85 y later.

In conclusion, it seems possible that large differences in ^2^H uptake between small/ juveniles and adult life stages of Malacostraca is related to molt-intermolt periods, but more detailed incubation studies with ^2^H and Malacostraca are required to confirm this.

### Metabolic activity and turnover of macrozoobenthos

An experiment aiming to determine *MTR* should include a sufficiently large number of replicates of the target taxon to allow sampling of multiple specimens (min *n* = 3) at T_1_, and potentially at T_2_, T_3_, etc. This is relatively easy to achieve when the target taxon is a species that can be kept *ex-situ* in an aquarium over longer periods of time. However, when the taxon is more delicate, such as sponges that can be sensitive to water flow (Hummel et al., 1988) and seawater composition (Maldonado et al., 2021), *ex-situ* and *in-situ* experiments may have to be combined. Indeed, here we coupled a short-term *ex-situ* incubation experiment (‘incubation’ or ‘pulse’ period: 12 h; (Stratmann et al., 2026)) with a long-term *in-situ* incubation experiment (this study), in which the taxa were kept *in-situ* for a 14 d ‘turnover’ or ‘chase’ period (‘incubation’/ ‘pulse’ period: 9 h), assuming that the taxa incorporated comparable amounts of ^2^H. The difference in the length of the ‘pulse’ periods is unfortunate and might lead to an overestimation of *MTR*. However, if this was the case, the overestimation would have affected all studied taxa equally. Consequently, this study might not have the best study design to use it as a comparison for future experiments because of this inconsistency, but at least it allows us to compare *MTR* among different taxa present in this study.

The highest *MTR* were measured in *H. panicea* and in *M. gigas*, whereby both macrozoobenthos species had higher *MTR* in summer (*H. panicea*: 4.76 % d^-1^; *M. gigas*: 2.28 d^-1^) than in autumn 2020 (*H. panicea*: 3.82 % d^-1^; *M. gigas*: 1.72 d^-1^; Fig. 9). This is intriguing as it shows that we were able to detect seasonal and taxonomic difference in *MTR*, though it is not uncommon to discover seasonal variation in benthic metabolic rates. For instance, at Fællestrand bight, island of Fyn (Danish Straits/ Belt Sea, Denmark) (Kristensen, 1993) could link changes in primary production to seasonal fluctuations in water temperature and daylight, whereas dynamics in benthic respiration were strongly linked to water temperature. In comparison, at Rothera Point, Antarctica, seasonal variations in metabolic rate of the shallow-water sea urchin *Sterechinus neumayeri* were mostly the result of feeding, growth and spawning activity, while water temperature had only a minor effect (Brockington and Clarke, 2001).

To our surprise, *MTR*s of *C. maenas, E. sinensis, P. depurator*, and Pycnogonida followed an opposite seasonal trend with higher *MTR* in autumn than in summer. This trend seems to contradict the metabolic theory of ecology (MTE) which postulates that metabolic activity is governed by a combination of temperature and body size (Brown et al., 2004) and increases with rising temperature. However, in August 2020 the daily mean water temperatures in the Eastern Scheldt exceeded 20ºC during a 13 day-long atmospheric heatwave (Hamer et al., 2026). During this heatwave, the intertidal fauna was exposed to above-optimal and partly sublethal water and air temperatures for several hours d^-1^ (Hamer et al., 2026). It is therefore possible that *C. maenas, E. sinensis, P. depurator*, and Pycnogonida were heat stressed and reduced their MTR, while *H. panicea* and *M. gigas* were better adapted to the extreme water and air temperatures. Hence, more experimental studies are required to assess the link between environmental fluctuations, such as temperature, and *MTR* before this approach can be used to test whether an ecosystem exposed to anthropogenic stressors is recovering.

## Conclusion

In this study, we investigated the long-term processing of ^13^C- and ^15^N-enriched bacterioplankton by an intertidal mudflat and Pacific oyster reef community in summer and autumn 2020. Fourteen days after the *in-situ* incubation experiments, most of the bacterioplankton-derived ^13^C and ^15^N was still detectable in sessile filter and suspension feeders, like sponges (*H. panicea, H. perlevis*), limpets (*C. fornicata*), and clams (*R. philippinarum*).

Additionally, we studied whether ^2^H uptake as a proxy for metabolic activity varied between life stages and detected partly significant differences in Malacostraca after 14 d. Whereas juvenile/ small *R. harrisii* had taken up almost twice as much ^2^H compared to the adult conspecifics, the ^2^H incorporation of adult *C. maenas* was almost the double of juvenile/ small *C. maenas*. For both cases, we proposed a link between ^2^H uptake and molt-intermolt periods in Malacostraca. Furthermore, we measured seasonal and taxonomic differences in *MTR* with the highest rates being measured in *H. panicea* and in *M. gigas* in summer and autumn 2020, and in Pycnogonida in autumn 2020. Due to a potential influence of temperature and other environmental factors on *MTR*, further studies are necessary before this method is suitable to investigate ecosystem recovery.

## Acknowledgments

We acknowledge the assistance of Pieter van Rijkswijk, Jeroen van Dalen, Anton Tramper, Jim van Belzen (all NIOZ-EDS), and Sara Stratmann while setting-up and conducting the experiments. We thank Jan Pene and Peter van Breugel (both NIOZ-EDS), Jort Ossebar, and Ronald van Bommel (both NIOZ-MMB) for their technical support during sample processing.

This research was funded by the JPI Oceans – Ecological Aspects of Deep-Sea Mining projects under NWO-ALW grant 856.18.003 and under NWO grant NWA.1745.23.1. TS was further supported by the Dutch Research Council NWO (NWO-Rubicon grant no. 019.182 EN.012, NWO-Talent program Veni grant no. VI. Veni.212.211, NWO Open Competition Domain Science – XS grant no. OCENW.XS24.2.193) and by the European Research Council (ERC-StG grant no. 101221692, project SPYCLING).

